# Evaluation of yield, yield stability and yield-protein trade-off in commercial faba bean cultivars

**DOI:** 10.1101/843862

**Authors:** Cathrine Kiel Skovbjerg, Jens Nørgaard Knudsen, Winnie Füchtbauer, Jens Stougaard, Frederick L. Stoddard, Luc Janss, Stig Uggerhøj Andersen

**Affiliations:** Department of Molecular Biology and Genetics, Aarhus University, DK-8000 Aarhus, Denmark; Nordic Seed, DK-8300 Odder, Denmark; Sejet Planteforædling, DK-8700 Horsens, Denmark; Department of Agricultural Sciences, Viikki Plant Science Centre and Helsinki Sustainability Centre, University of Helsinki, FI-00014 Helsinki, Finland

**Keywords:** *Vicia faba*, yield, yield stability, yield-protein trade-off, potential crop, genotype x environment interaction

## Abstract

Faba bean is a legume crop with high protein content and large potential for cultivation in the Northern latitudes. However, it has a reputation for being an unstable crop with large inter-annual variability, mostly explained by yearly variation in rainfall. Consequently, the objective is to breed cultivars with high seed yield and high yield stability. In this study, 17 commercial cultivars of faba bean were evaluated for seed yield, yield stability and trade-off between seed yield and protein content in four locations in Denmark and Finland during 2016-2018. We found that location and year effects accounted for 72% of the total seed yield variation. Cultivar by environment interactions (G×E) were found to be small and did not cause re-ranking of cultivars in different environments. Yield stability contributed little to the mean yield of the cultivars because high-yielding cultivars consistently outperformed the lower yielding genotypes, even under the most adverse conditions. The latter was also the case for total protein yield quantified as total yield multiplied by seed protein percentage. Although we found a strong negative correlation of −0.64 between yield and protein content, a few cultivars produced high yields while maintaining a relatively high protein content, suggesting that these traits may to some degree be genetically separable.

## 1. Introduction

Faba bean (*Vicia faba* L.) is an annual legume with a high content of protein and the ability to fix atmospheric N_2_ in symbiosis with compatible rhizobia. The nitrogen fixing ability makes it a good crop for sustainable agricultural systems as the need for N fertilizer input is minimal. Moreover, the high protein content of seeds makes it a good source of both animal feed and human food (Stoddard et al., 2009; Crépon et al., 2010). According to FAOSTAT (2019) the largest producers of faba bean are China, Ethiopia and Australia, that together account for almost 65% of the production worldwide. Currently, the production of faba bean in northern Europe accounts for less than 5% of the total production (FAOSTAT 2019), and to meet the need for supplementary protein in animal feed, European countries import large amounts of soybean (De Visser et al., 2014).

Faba bean is cold-hardy and grows best under cool and moist conditions making it a very attractive legume crop for northern countries (Duc et al., 2015). It has shown good potential for use in pig feed, and is associated with higher protein content than other common food legumes such as peas (Partanen et al., 2006). Replacing part of the soybean import in European countries with locally grown faba bean will not only benefit the environment but also the national economies (Stoddard et al., 2009).

However, the yield of faba bean, like that of many other legumes, is considered unstable because of large inter-annual variability in yield compared to non-legume species (Cernay, 2015). Much of this instability is attributable to the contrast between spring sowing of most legume crops and autumn sowing of most of the cereals with which they are compared (Reckling et al., 2018a). Faba bean is considered drought susceptible, and variation in the amount of rainfall has been reported as a major cause of seed yield instability (Link et al., 1999). To make faba bean a more attractive crop to agronomists and farmers, the identification of cultivars with high seed yield and high yield stability is desirable. When collecting data from multi-environment yield trials, yield stability can be investigated as the ability of some genotypes to perform consistently over a wide range of environments (Finlay and Wilkinson, 1963). The demands for cultivars better adapted to changing conditions is of great interest even in what seems to be consistent environments, as climate conditions change from year to year and are expected to fluctuate more in the future as a consequence of global warming. There is already an indication that grain legume yield variability has increased in the last 60 years (Reckling et al., 2018b).

Several methods may be used to assess yield stability. Static (type 1) yield stability refers to genotypes that perform consistently in all environments, i.e., environmental stability, whereas dynamic (type 2) yield stability refers to genotypes that show no genotype x environment (G×E) interaction (Annicchiarico, 2002). Among the most common measures of static yield stability are the environmental variance and coefficient of variation, whereas the Finlay-Wilkinson regression, which works by regressing performance of each genotype on the environmental means, is a popular method used by plant breeders to describe the G×E interaction. G×E can be classified as G×E that causes re-ranking of cultivars between environments, and a milder type of G×E that does not cause re-reranking but where differences between cultivars are larger in some environments than in others. Covering both the static and the dynamic types of stability is expected to benefit the identification of cultivars with stable yield.

The objectives of this study were to evaluate the seed yield of 17 commercial cultivars of faba bean across different Nordic environments, to determine the stability of yield for each cultivar and to assess the possible trade-off between seed yield and protein content of seeds.

## 2. Materials and methods

### 2.1 Plant material and growing conditions

Seventeen commercial spring-type cultivars of faba bean originating from cool climates (Table 1) were grown in four Nordic locations during 2016-2018. Three of the locations were in Denmark: Dyngby (55.942°N, 10.208°E), Nørre Aaby (55.470°N, 9.845°E) and Horsens (55.830°N, 9.950°E). The fourth location was the University of Helsinki research farm at Viikki, Helsinki, Finland (60.224°N, 25.021°E). The combination of a year and location is considered as an environment. The trials were sown in April to May and harvested in August to September. The fields in Denmark and Finland were set up with three and four replicates of each cultivar, respectively. The position of each plot was scored so the effects of the field position and neighbor phenotypes could be included in the data analyses. Plot sizes were 5.5 m × 1.5 m in Dyngby, 7.0 m × 1.5 m in Horsens and Nørre Aaby and 9.0 m × 1.25 m in Viikki.

**Table 1.**
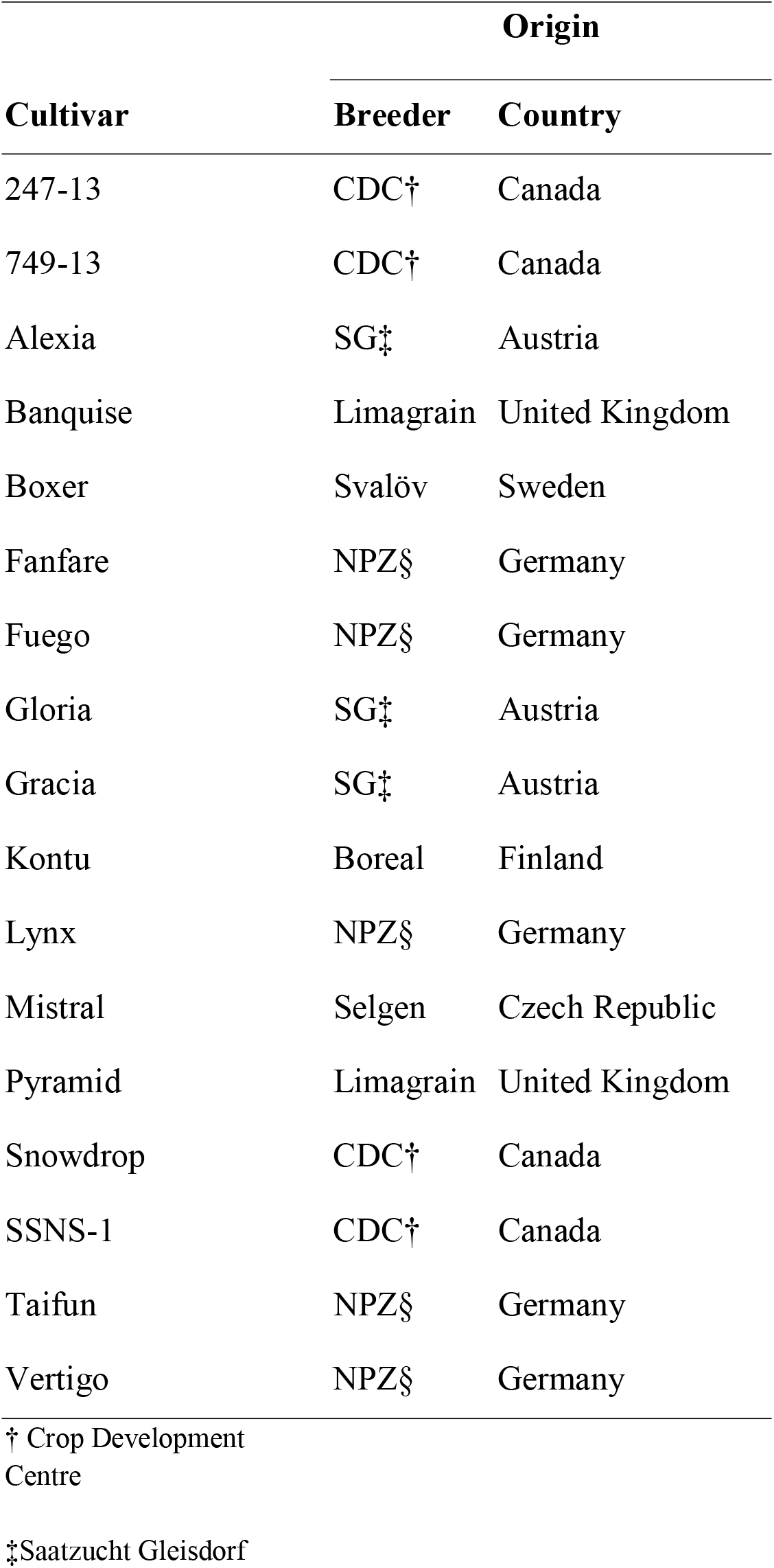
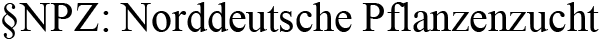
Overview of cultivar origin

| Cultivar | Origin |  |
| --- | --- | --- |
|  | Breeder | Country |
| 247-13 | CDC† | Canada |
| 749-13 | CDC† | Canada |
| Alexia | SG‡ | Austria |
| Banquise | Limagrain | United Kingdom |
| Boxer | Svalöv | Sweden |
| Fanfare | NPZ§ | Germany |
| Fuego | NPZ§ | Germany |
| Gloria | SG‡ | Austria |
| Gracia | SG‡ | Austria |
| Kontu | Boreal | Finland |
| Lynx | NPZ§ | Germany |
| Mistral | Selgen | Czech Republic |
| Pyramid | Limagrain | United Kingdom |
| Snowdrop | CDC† | Canada |
| SSNS-1 | CDC† | Canada |
| Taifun | NPZ§ | Germany |
| Vertigo | NPZ§ | Germany |

† Crop Development Centre
‡Saatzucht Gleisdorf
§NPZ: Norddeutsche Pflanzenzucht

No plots were evaluated for yield in Horsens and Nørre Aaby during 2016, due to a treatment mistake, and no harvest took place in Viikki during 2017 due to autumn weather damage, so there was a total of 9 environments for yield data. Plots were harvested with plot-scale combine harvesters.

### 2.2 Phenotyping

Seed moisture content (%) and protein content (%) were determined with a NIR sensor (DA 7440, Perten, Stockholm, Sweden or Infratec 1241, FOSS, Hillerød, Denmark) on samples from Dyngby in all years and Horsens in 2017. For samples from Finland in 2016, protein content was determined using the Kjeldahl method. After seed cleaning, plot yields (g/m^2^) and protein contents were corrected to 15% moisture content. This gave a total yield data set of 487 observations and a protein data set of 270 observations.

### 2.3 Statistical analyses

#### 2.3.1 Yield analyses

For the analysis of yield measured across different years and locations the following linear mixed model was applied. The mixed model was fit in R (v. 3.5.1) (R Core Team, 2018) using the ‘lme4’ package (Bates, 2015).

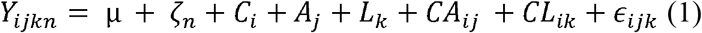

Where Y_ijkn_ is a random variable representing the yield (g/m^2^) of a plot, C_i_ is the effect of the ith cultivar (i=1,2,…17), A_j_ is the effect of the jth year (j=1,2,3), L_k_ is the effect of the kth location (k=1,2,…4), CA_ij_ is the effect resulting from cultivar × year interaction, CL_ik_ is the effect resulting from cultivar × location interaction, ε_ijk_ is the residual error term of the ith cultivar in the jth year and kth location, μ is the overall mean and ζ_n_ is a fixed effect describing the average yield of the neighbor plots.

The model assumes normality of data and independence of residuals. Visual inspection of QQ-plots and plots checking for homoscedasticity revealed no violations of assumptions. This was further confirmed by a Shapiro-Wilk test of normality of residuals yielding a p-value of 0.11, meaning no significant evidence of the data deviating from normality on a 0.05 significance level.

From the model, restricted maximum likelihood (REML) estimates of different variance components were extracted as well as estimates of fixed effects. The variance components were used to calculate proportions of the total phenotypic variance explained by single components. In addition, estimates of variance were used for calculation of differently defined broad-sense heritability terms, including heritability of a single plots, heritability of cultivar means and heritability of cultivar means within a single year and location. The equations used to obtain the heritability estimates were the following:

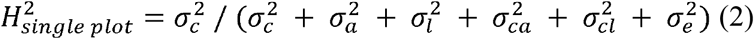

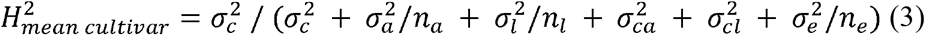

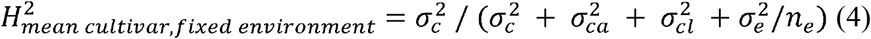

Where 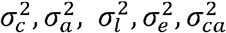 and 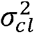 are the variances of the cultivar, year, location, residual, cultivar with year interaction and cultivar with location interaction, respectively; *n*_*a*_ denotes the number of years, *n*_*l*_ denotes the number of locations and *n*_*e*_ denotes the average number of replications of each cultivar in the data set. In principle, the broad-sense heritability of a cultivar mean can become 1 if the number of plots in the experiments becomes very large.

Cultivar best linear unbiased predictors (BLUPs) were extracted from the model (1) and used to evaluate the effect of the different cultivars on yield. P-values associated with fixed effects were extracted using the R-package “afex” (Singmann, 2019)

#### 2.3.2 Evaluation of yield stability

To examine the yield stability of the different cultivars across different environments, three different yield stability parameters were included in the analyses: Finlay-Wilkinson regression coefficient, coefficient of variation (CV) and the variance of the logarithmic yield data set.

##### 2.3.2.1 Finlay-Wilkinson regression coefficient

Yield stability was accessed across different environments by the Finlay-Wilkinson regression coefficient (Finlay and Wilkinson, 1963). The yield response of a given cultivar can be represented as:

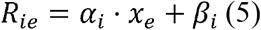

Where R_ie_ is the modeled yield response of the ith cultivar in the eth environment, *β*_*i*_ is the intercept value of ith cultivar, and x_e_ is the mean yield in the eth environment, *α*_*i*_ is the regression coefficient of the ith cultivar and is a measure of genotype by environment interaction. The lower the regression coefficient, the greater resistance of the cultivar to environmental change, i.e., the higher its stability.

##### 2.3.2.2 Coefficient of variation

The coefficient of variation (CV) measures the variation of a cultivar across environments. It can be defined as:

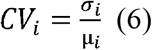

Where *σ*_*i*_ and μ_*i*_ is the standard deviation and mean of yield data for the ith cultivar, respectively. Cultivars with the highest stability will give rise to the lowest coefficients of variations.

##### 2.3.2.3 Variance of logarithmic yield within cultivar

As another measure of yield stability, we calculated the variance of the logarithm of yield of each cultivar across environment. The logarithmic scale was included to account for the fact that yield variances are dependent on yield levels.

#### 2.3.3 Genetic correlations

Genotypic correlation coefficients between seed yield and protein content of seeds were estimated as the Pearson correlation between the sets of cultivar BLUPs obtained by using protein or yield as the random variable in the model (1). Pearson correlations were calculated using the “correlation” function in the “agricolae” R-package (de Mendiburu, 2010).

The R-script used for statistical analyses can be found at https://github.com/cks2903/Faba_bean_yield_2019

## 3. Results

### 3.1 Yield and yield components

The average, maximum and minimum yield of the 17 commercial faba bean cultivars are listed in Table 2. Average cultivar yields varied from 296.4 g/m^2^ in Kontu to 490.1 g/m^2^ in Lynx. The minimum yield was observed in Kontu as 86.9 g/m^2^, and the maximum yield of 962.4 g/m^2^ was observed in Fanfare. Minimum cultivar yields were all below 200 g/m^2^ whereas the interval of the maximum cultivar yields was larger (489 to 962.4 g/m^2^).

**Table 2.** Average, maximum and minimum yield and protein content of seeds of the 17 commercial faba bean cultivars

| **Cultivar** | **Seed yield (g/m<sup>2</sup>)** |  |  | **Protein content of seeds (%)** |  |  | **Protein yield (g/m<sup>2</sup>)** |
| --- | --- | --- | --- | --- | --- | --- | --- |
|  | **Avg.** | **Max.** | **Min.** | **Avg.** | **Max.** | **Min.** | **Avg.** |
| 247-13 | 410.8 | 820.6 | 137.1 | 29.3 | 31.9 | 27.7 | 151.7 |
| 749-13 | 449.5 | 783.0 | 137.1 | 28.3 | 29.6 | 26.9 | 149.9 |
| Alexia | 429.6 | 800.0 | 145.3 | 29.2 | 30.9 | 27.4 | 157.1 |
| Banquise | 450.7 | 833.9 | 164.8 | 28.6 | 31.0 | 27.2 | 152.5 |
| Boxer | 455.0 | 901.8 | 167.1 | 28.4 | 30.4 | 26.9 | 161.2 |
| Fanfare | 484.4 | 962.4 | 168.5 | 28.7 | 30.9 | 27.5 | 175.2 |
| Fuego | 460.3 | 838.8 | 187.6 | 28.7 | 31.6 | 26.7 | 164.7 |
| Gloria | 396.5 | 787.9 | 151.9 | 31.9 | 34.3 | 29.5 | 161.5 |
| Gracia | 451.0 | 916.4 | 154.3 | 29.4 | 31.8 | 26.7 | 168.3 |
| Kontu | 296.4 | 586.7 | 86.9 | 30.6 | 33.5 | 28.8 | 113.3 |
| Lynx | 490.1 | 864.2 | 198.1 | 28.8 | 30.0 | 26.6 | 176.7 |
| Mistral | 408.4 | 722.4 | 168.6 | 31.6 | 34.9 | 28.8 | 158.3 |
| Pyramid | 481.5 | 938.2 | 156.5 | 28.4 | 31.4 | 26.9 | 169.9 |
| Snowdrop | 388.9 | 489.0 | 122.9 | 30.3 | 32.4 | 28.4 | 123.2 |
| SSNS-1 | 388.9 | 770.9 | 150.5 | 30.7 | 33.6 | 27.4 | 146.7 |
| Taifun | 448.9 | 807.3 | 168.4 | 28.7 | 30.0 | 26.7 | 160.1 |
| Vertigo | 476.5 | 947.9 | 159.5 | 29.3 | 31.6 | 27.1 | 176.1 |

Abbreviations: Avg, average; Max, maximum; Min, minimum.

Estimates of cultivar, year, location, cultivar × location interaction, cultivar × year interaction, residual variances and the proportion explained by the different components are presented in Table 3. The largest proportions of the variance of seed yield was explained by year (53.8%) and location (18.2%), together accounting for 72.0% of the total phenotypic variance in yield. This indicates a large environmental effect on seed yield, which was further supported by G×E interaction effects accounting for 5.7% of the total variance. The cultivar was estimated to account for 11.2% of the variance, indicating that main-effects of genotypes account for a larger proportion of variance than G×E interactions. The average neighbor score was estimated to have a significant effect of 0.40 ± 0.04 (P < 2e−16).

**Table 3.** Summary of variance components of joint yield analysis on the 17 cultivars in 9 environments

| Source | Variance component | Variance proportion (%) |
| --- | --- | --- |
| Cultivar | 2605.5 | 11.2 |
| Cultivar x location | 446.6 | 1.9 |
| Cultivar x year | 880.7 | 3.8 |
| Year | 12506.1 | 53.8 |
| Location | 4240.2 | 18.2 |
| Residual | 2555.0 | 11.0 |

The estimated broad sense heritability on a single plot and the mean cultivar was 0.11 and 0.28, respectively. As year and location explain the largest proportion of the variance, the broad-sense heritability of the cultivar mean within a given environment was estimated to be 0.84.

### 3.2 Yield stability

To compare different parameters of stability, yield stability was evaluated as variance of logarithmic yield, coefficient of variation (CV%) and Finlay-Wilkinson regression coefficients. The results are summarized in Table 4. Variance of logarithmic yield varied from 191.8 * 10^−3^ (Fuego) to 262.9 * 10^−3^ (Gloria) and had a mean of 217.1 * 10^−3^. Coefficient of variation (CV%) varied from 36.1% (Snowdrop) to 47.5% (Gloria) with a mean of 41.7%. Finlay-Wilkinson regression coefficients varied from 0.60 (Snowdrop) to 1.15 (Fanfare).

**Table 4.**
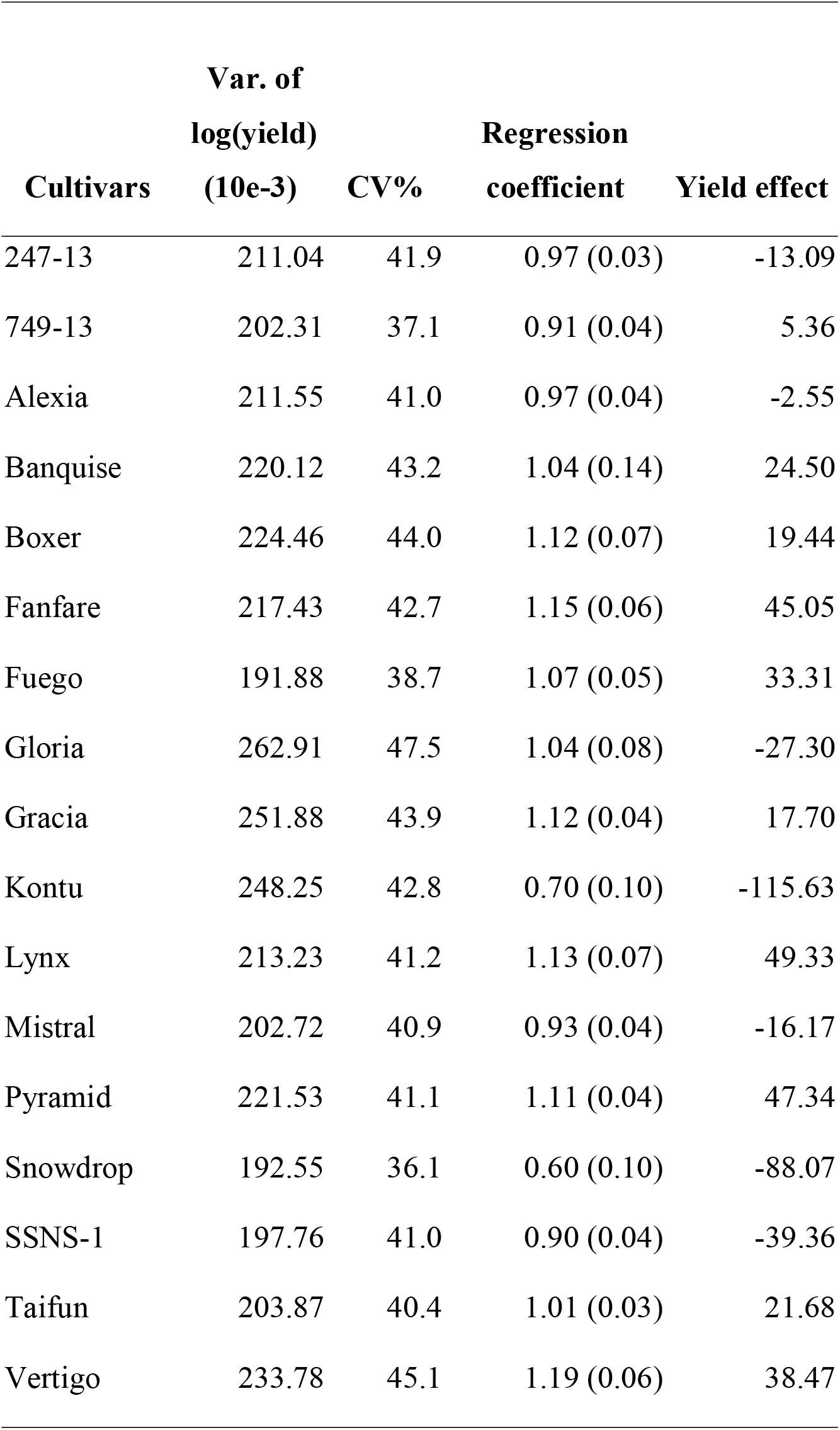
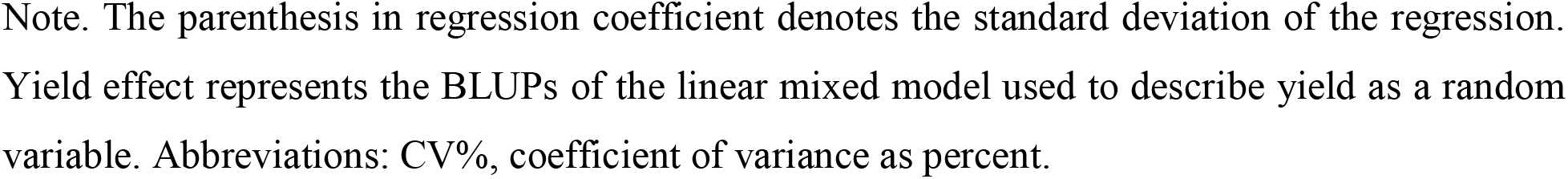
Yield stability in the 17 faba bean cultivars

| **Cultivars** | **Var. of**<br>**log(yield)**<br>**(10e-3)** | **CV%** | **Regression**<br>**coefficient** | **Yield effect** |
| --- | --- | --- | --- | --- |
| 247-13 | 211.04 | 41.9 | 0.97 (0.03) | -13.09 |
| 749-13 | 202.31 | 37.1 | 0.91 (0.04) | 5.36 |
| Alexia | 211.55 | 41.0 | 0.97 (0.04) | -2.55 |
| Banquise | 220.12 | 43.2 | 1.04 (0.14) | 24.50 |
| Boxer | 224.46 | 44.0 | 1.12 (0.07) | 19.44 |
| Fanfare | 217.43 | 42.7 | 1.15 (0.06) | 45.05 |
| Fuego | 191.88 | 38.7 | 1.07 (0.05) | 33.31 |
| Gloria | 262.91 | 47.5 | 1.04 (0.08) | -27.30 |
| Gracia | 251.88 | 43.9 | 1.12 (0.04) | 17.70 |
| Kontu | 248.25 | 42.8 | 0.70 (0.10) | -115.63 |
| Lynx | 213.23 | 41.2 | 1.13 (0.07) | 49.33 |
| Mistral | 202.72 | 40.9 | 0.93 (0.04) | -16.17 |
| Pyramid | 221.53 | 41.1 | 1.11 (0.04) | 47.34 |
| Snowdrop | 192.55 | 36.1 | 0.60 (0.10) | -88.07 |
| SSNS-1 | 197.76 | 41.0 | 0.90 (0.04) | -39.36 |
| Taifun | 203.87 | 40.4 | 1.01 (0.03) | 21.68 |
| Vertigo | 233.78 | 45.1 | 1.19 (0.06) | 38.47 |

Note. The parenthesis in regression coefficient denotes the standard deviation of the regression. Yield effect represents the BLUPs of the linear mixed model used to describe yield as a random variable. Abbreviations: CV%, coefficient of variance as percent.

Plotting the variance of the logarithmic yield of each cultivar against its mean yield allows us to identify high-yielding cultivars with low yield variance i.e. high stability (Figure 1A). Cultivars that showed yield above average, that is a positive yield effect in Table 4, but low variance of logarithmic yield were 749-13, Fanfare, Fuego, Lynx and Taifun.

**Figure 1.**
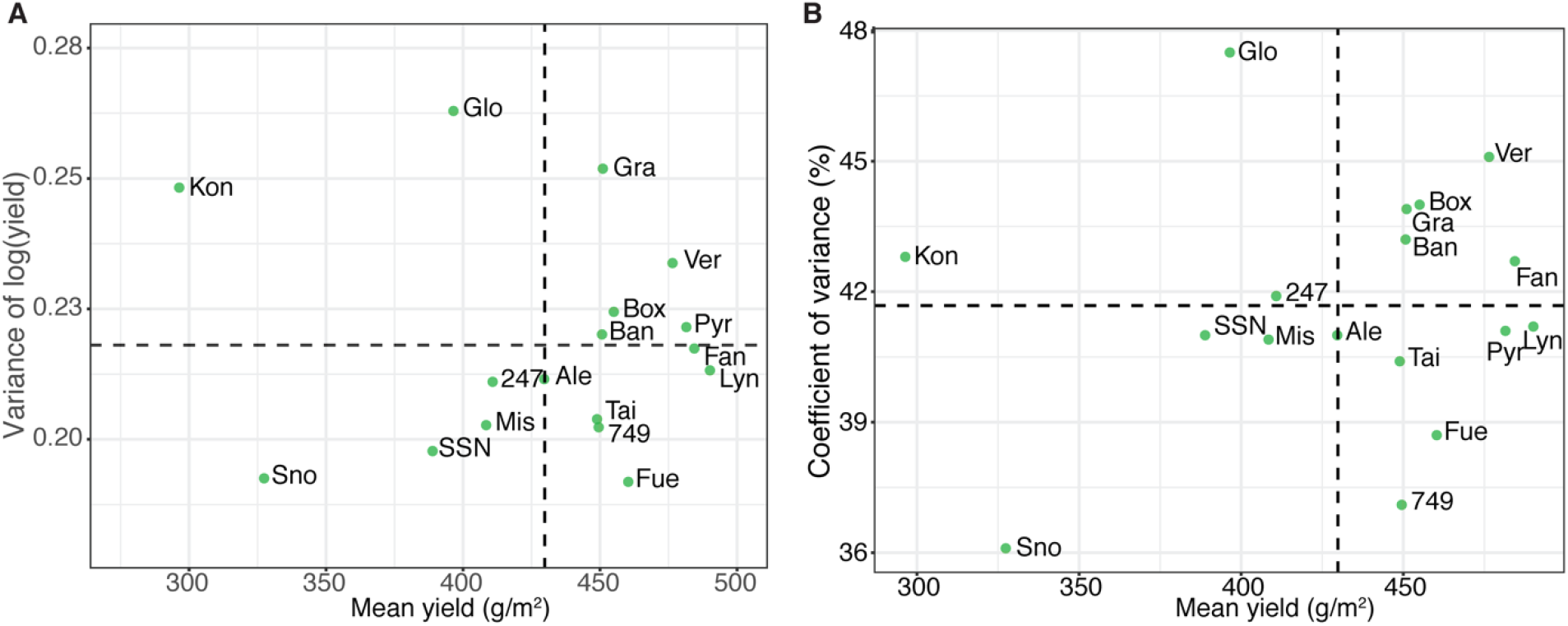
Mean seed yield plotted against the variance of the logarithmic yield (A) or the coefficient of variance in percent (CV%) (B) of the 17 cultivars. Dashed lines represent the average yield of all cultivars (vertical) and the average of the yield stability parameter across all cultivars (horizontal). Abbreviations. 247: 247-13, 749: 749-13, Ale: Alexia, Ban: Banquise, Box: Boxer, Fan: Fanfare, Fue: Fuego, Glo: Gloria, Gra: Gracia, Kon: Kontu, Lyn: Lynx, Mis: Mistral, Pyr: Pyramid, Sno: Snowdrop, SSN: SSNS-1, Tai: Taifun, Ver: Vertigo.

A similar plot was made for yield stability measured as coefficient of variation (CV%) (Figure 1B). From the data presented in Figure 1B and Table 4 we identified cultivars 749-13, Fuego, Lynx, Pyramid and Taifun cultivars to have high yield and low CV%.

To evaluate the G×E interaction in different cultivars, the mean yield of a cultivar in the given environment was plotted against the year-location mean yield and regression was performed. Figure 2 shows the regressions of the 17 faba bean cultivars. The regression coefficients are listed in Table 4 as well. Regression coefficient above 1 describes genotypes that are more sensitive to the environment than the average, whereas regression coefficients below 1 describes genotypes less sensitive to the environment than average.

**Figure 2.**
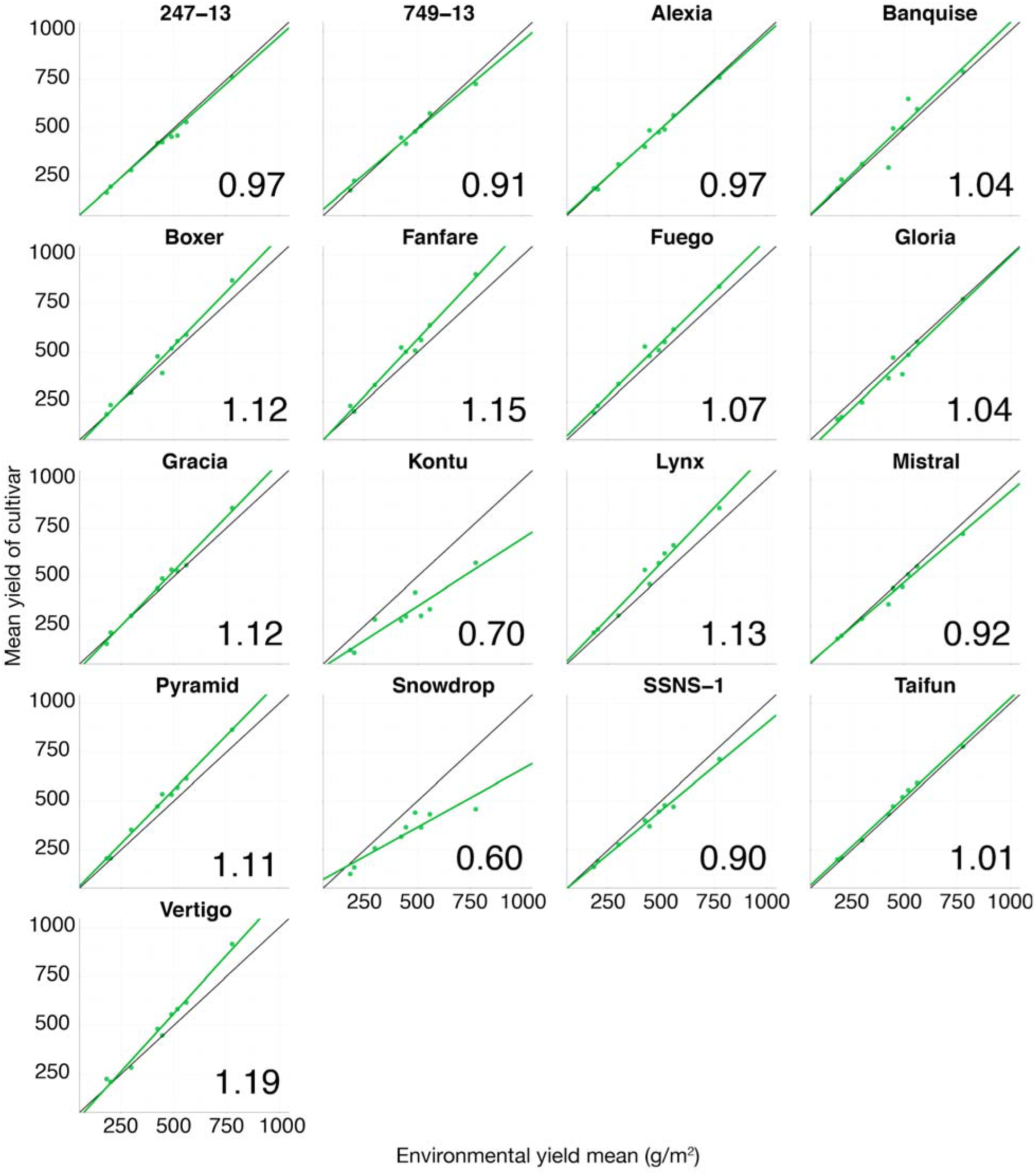
Regression of mean seed yield of a cultivar in a given environment against the environmental mean yields. The black lines display the average G×E effect. Numbers in the lower corner refer to the regression coefficient of the fitted line (green).

Regression coefficients indicated that cultivars 749-13, Kontu, Mistral, Snowdrop and SSNS-1 were more resistant to environmental changes than average. Especially low-yielding cultivars Snowdrop and Kontu showed low G×E interaction.

The high yielding cultivars Fuego and Taifun seemed to have regressions coefficients close to 1, i.e. average G×E interaction, but performed better than or similar to the environmental means in all environments, in contrast to cultivar 749-13 which performed worse than the environmental means in some environments. High-yielding cultivars Fanfare, Lynx and Pyramid performed better than the average in all environments but showed the largest G×E interaction terms. G×E caused no re-ranking of cultivars in different environments. The correlation between the Finlay-Wilkinson regression coefficients and the average seed yield of cultivars was found to be 0.90 (P = 6.15 * 10^−7^).

### 3.3 Protein-yield trade-off

The average, maximum and minimum protein content of seed of the 17 commercial faba bean cultivars are listed in Table 2. The average protein content of seeds varied from 28.3% in 749-13 to 31.9% in Gloria. The minimum protein content was observed in Lynx (26.6%) and the maximum protein content was observed in Mistral (34.9%).

To investigate if there was a trade-off between seed yield and protein content of seeds in the 17 commercial cultivars of faba bean, Pearson correlation coefficients were calculated between the estimations of random effects associated with each cultivar in the mixed model, i.e. the best linear unbiased predictors (BLUPs) obtained for the traits individually. Protein content of seeds and seed yield showed a high negative genetic correlation of −0.64 (P = 0.0053). In Figure 3 cultivar BLUPs obtained when predicting yield were plotted against the BLUPs obtained when predicting protein content, and a linear model was fitted. Most cultivars follow the regression showing a decreasing effect on yield, as the effect on protein increases. Cultivars that deviate from the regression line are of particular interest, especially those above the line as such cultivars will be desirable having both high yield and high protein content. Considering the 95% confidence interval of the regression line (the shaded area), the cultivars Gloria, Gracia, Mistral, SSNS-1 and Vertigo showed higher protein content than would be expected by their yield (Figure 3). The cultivar effects of Vertigo and Gracia were found to be positive for both protein content and seed yield. Cultivars 247-13, 749-13, Alexia, Banquise and Boxer also deviate from the line, but all show a lower protein content than would be expected from their seed yield (Figure 3). High yielding/high stability cultivars Fuego and Taifun did not deviate from the regression line, indicating a large protein trade-off for these cultivars.

**Figure 3.**
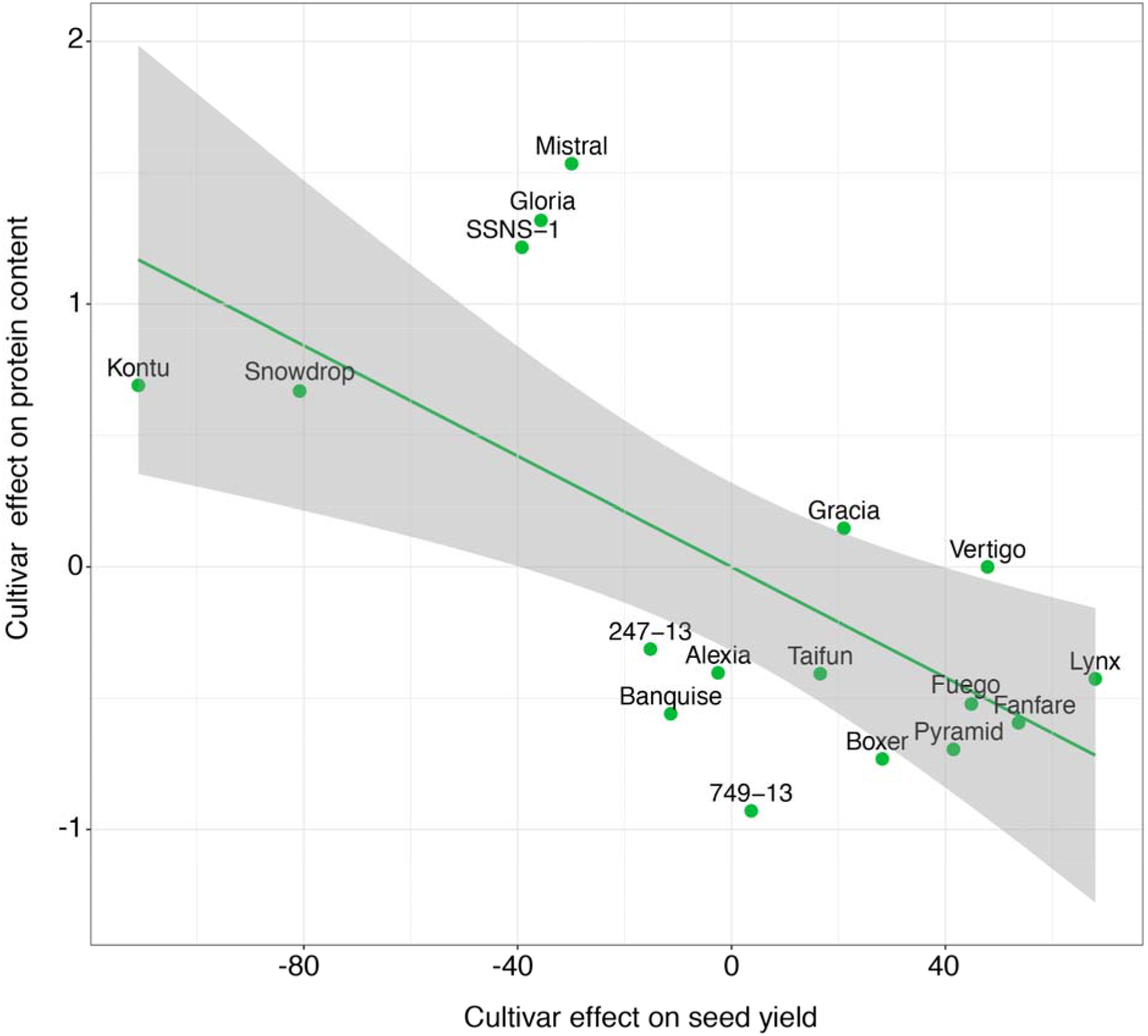
Genetic correlation between the protein content of seeds and the seed yield of the 17 commercial faba bean cultivars. The shaded area specifies the 95% confidence interval of the regression line.

### 3.4 Protein yield stability

To take the large negative genetic correlation of protein and yield into account, when considering trait stability, similar plots to Figure 2 were produced for protein yield (g/m^2^) (Figure 4). Protein yield was defined as the weight of protein that was harvested from seeds in a plot. It was calculated by multiplying protein content of seeds with seed yield (g/m^2^) (Table 5). For stability of protein yield cultivar Fuego and Pyramid had regression coefficients below 1 but performed better than or equal to the average environmental protein yields in all environments, indicating a lower G×E interaction for the protein yield trait than for yield alone. Taifun had a regression coefficient below 1, but showed lower than average protein yield in high-yielding environments (Figure 2, Figure 4). The high yielding cultivar Lynx had a regression coefficient close to 1 (1.05), i.e. average G×E interaction on the protein yield trait, whereas it showed one of the largest G×E interactions on the yield trait alone. For both traits Lynx outperformed the means in all environments investigated. Fanfare showed an above-average G×E interaction for protein yield (1.14) but did not outperform the average phenotype in all environments as we observed for seed yield. Gracia performed better than the mean protein yield in every environment, whereas it performed below the environmental mean in low-yielding environments when looking at seed yield alone (Figure 2, Figure 4).

**Table 5.** Protein yield stability in the 17 faba bean cultivars

| **Cultivars** | **Regression**<br>**coefficient** |
| --- | --- |
| 247-13 | 1.03 (0.10) |
| 749-13 | 0.84 (0.08) |
| Alexia | 1.01 (0.14) |
| Banquise | 1.08 (0.29) |
| Boxer | 1.19 (0.16) |
| Fanfare | 1.10 (0.07) |
| Fuego | 0.95 (0.15) |
| Gloria | 1.13 (0.19) |
| Gracia | 1.11 (0.06) |
| Kontu | 0.87 (0.10) |
| Lynx | 1.05 (0.20) |
| Mistral | 1.01 (0.19) |
| Pyramid | 0.95 (0.12) |
| Snowdrop | 0.36 (0.17) |
| SSNS-1 | 0.96 (0.11) |
| Taifun | 0.94 (0.04) |
| Vertigo | 1.29 (0.11) |

Note. The parenthesis in regression coefficient denotes the standard deviation of the regression.

**Figure 4.**
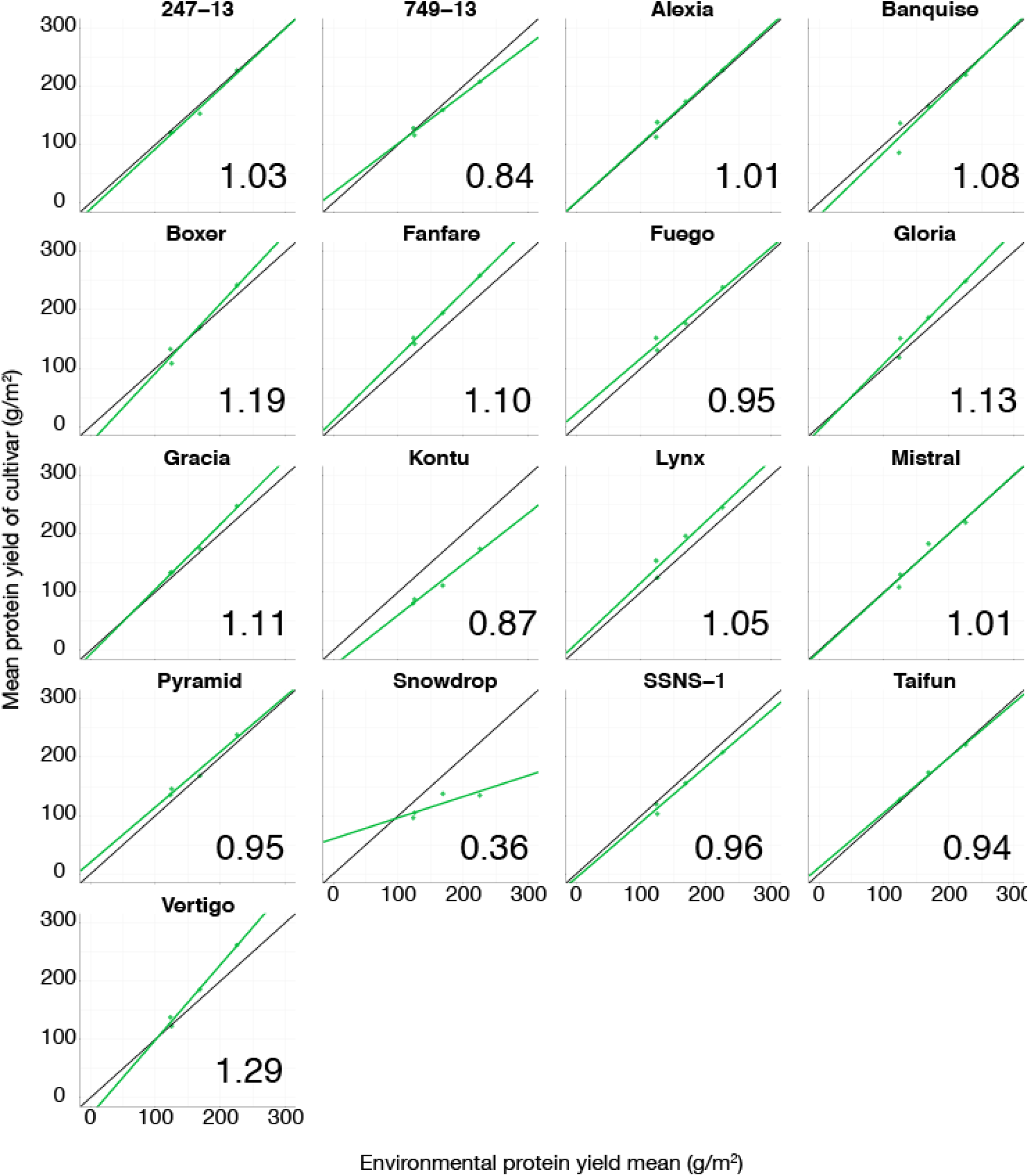
Regression of mean protein yield of a cultivar in a given environment against the environmental mean protein yields. The black lines display the average G×E effect. Numbers in the lower corner refer to the regression coefficient of the fitted line (green).

## 4. Discussion

Our results show that there is variance in seed yield and yield stability of the 17 commercial faba bean cultivars evaluated in this study. There is a tendency of an inverse relationship between yield and yield-stability, with low-yielding cultivars showing the highest stability and higher-yielding cultivars showing lower stability. This tendency has also been reported in previous evaluations of yield stability in different faba bean genotypes (Temesgen, 2015). The environments evaluated in this study showed pronounced variation in mean seed yield ranging from 182.9 g/m^2^ to 776.1 g/m^2^ in Horsens, 2018 and Dyngby, 2017, respectively. Because the environments evaluated here represent a large range of Nordic environments, cultivars that performed better than the mean in every environment are expected to generally perform well across most Nordic environments used for faba bean cultivation. This indicates that cultivating high-yielding genotypes with a large G×E interaction does not negatively affect the mean seed yield, as these cultivars can in fact benefit from a large G×E interaction. However, for breeders large G×E interactions would limit response to selection if G×E is so substantial that different environments require development of specialized varieties for each environment. In the faba bean cultivars evaluated here, this is not the case as G×E does not cause reranking of cultivar performance across environments. In addition to the high-yielding cultivars, we found several cultivars that did not have the ability to exploit the high-yielding environments, consequently showing Finlay-Wilkinson regression coefficients below one.

We observed a negative genetic correlation between protein content of seed and seed yield, which agrees with findings of similar phenotypic correlations in pea (Jha, 2012), chickpea (Frimpong, 2009) and in recent studies of faba bean (Barłóg, 2019). Selecting for cultivars with high seed yield might to a certain degree mean selecting for low content of protein in seeds. However, a low content of protein in seeds can be overcome by higher seed yield, meaning altogether higher protein yield. In addition to the negative genetic correlation, a significant phenotypic correlation between seed yield and protein content of seeds were found (−0.14, P = 0.0248). The lower phenotypic correlation can be explained by year and locations having different effects on the two traits. The relationship between yield and protein are emphasized by the findings of low-yielding cultivars often being associated with high protein content of seeds and the other way around. However, we identified a few cultivars that produced high seed yields while maintaining a protein concentration above average. Another desired quality in faba bean cultivars is early maturation. However, there is a fundamental conflict between breeding cultivars with both early maturation and high seed yield, as the correlation between growing period and yield is typically found to be significantly positive (Prohens, J., Nuez, F., & Carena, M. J., 2008). Varieties bred to flower and mature very early such as Kontu will consequently be expected to show lower seed yield than varieties with a longer production time (Table 2).

The Finlay-Wilkinson regression coefficient used to describe yield stability was strongly correlated with seed yield (0.90, P = 6.15 * 10^−7^), whereas the other stability parameters of seed yield, variance of logarithmic yield and CV% were not significantly correlated with seed yield, showing Pearson correlation coefficients of −0.08 (P = 0.75) and 0.15 (P = 0.57), respectively. This indicates that using the regression coefficient to evaluate yield stability in agronomic breeding programs might favor development of cultivars with both high yield and high yield stability.

The correlation between the coefficient of variance and the variance of logarithmic yield was 0.84 (P = 2.62*10^−5^), the correlation between CV% and the Finlay-Wilkinson regression coefficient was 0.53 (P = 0.029) and the correlation between Finlay-Wilkinson regression coefficient and the variance of logarithmic yield was 0.26 (P = 0.32). The correlations between the different stability parameters reveal that each stability parameter describes different aspects of yield stability, thus contributing to an overall evaluation of yield stability.

The heritability of the cultivar mean yield was found to be relatively low (0.28) compared to what has earlier been reported by other researchers studying faba bean (0.62-0.77) (Toker, 2004, Alan 2007) or other legumes such as chickpeas (0.86) (Canci, 2009). The low heritability reflects the high environmental influence on seed yield, i.e. year and location effects explained 72% of all phenotypic variance in yield. In this study, the three years in the joint analysis were very different with 2018 being dry (50.9 mm (DK) or 81.5 mm (FN) precipitation from May to July) and 2017 being wet (204.2 mm (DK) precipitation from May to June), which gave rise to a large yield variance across years. The dataset used for yield evaluation in this study contained more observations from the dry year 2018 than from 2016 or 2017, and the broad-sense heritability of faba bean cultivars has previously been reported to be higher under well-watered conditions than under drought-conditions (Link,1999). In agreement with our findings, previous studies on seed yield has also found environmental effects to explain the largest proportion of phenotypic variance in faba beans (Temesgen, 2015) and chickpeas (Frimpong, 2009).

## 5. Conclusion

In this study we observed large variation in seed yield and yield stability of the 17 commercial faba bean cultivars evaluated. Seed yield was found to be a trait highly influenced by environment, i.e. the proportion of variance explained by environmental factors was larger than 72%.

We found that it was possible to identify high-yielding faba bean cultivars with high static yield stability. In addition, we found that G×E interactions caused no re-ranking of cultivars in different environments, and that high-yielding cultivars consistently outperformed the lower yielding genotypes in all environments, indicating that it could still be beneficial to grow cultivars with a high G×E interaction of seed yield since there is no trade-off of seed yield. A significant negative genetic correlation was observed between protein content of seed and seed yield. However, we identified a few cultivars which produced high yields while maintaining a relatively high protein content, suggesting that these traits may to some degree be genetically separable.

## Acknowledgments

The work was funded by NORFAB: Protein for the Northern Hemisphere 5158-00004B.

## Author contributions

Conceptualization, C.K.S., S.U.A., L.J.; Methodology, C.K.S., L.J.; Validation, C.K.S., L.J., S.U.A.; Investigation, J.N.K., W.F., F.L.S.; Formal Analysis, C.K.S.; Resources, J.N.K., W.F., F.L.S.; Data Curation, C.K.S., S.U.A.; Writing - Original Draft, C.K.S., S.U.A; Writing -Review & Editing, C.K.S., S.U.A., F.L.S, L.J.; Visualization, C.K.S.; Supervision, S.U.A., L.J.; Project Administration, S.U.A., J.S.; Funding Acquisition, S.U.A.,J.S..

## Data availability statement

The source data is available in supplementary file 1.

## References

Alan, O., & Geren, H. (2007). Evaluation of heritability and correlation for seed yield and yield components in faba bean (Vicia faba L.). Journal of Agronomy, 6, 484–487. Doi: 10.3923/ja.2007.484.487

Annicchiarico, P. (2002). Genotype × environment interactions: challenges and opportunities for plant breeding and cultivar recommendations. FAO Plant Production and Protection Paper No. 174. Rome, Italy: Food & Agriculture Organization of the United Nations. 115 pp.

Barłóg, P., Grzebisz, W., & Łukowiak, R. (2019). The Effect of Potassium and Sulfur Fertilization on Seed Quality of Faba Bean (*Vicia faba* L.). Agronomy, 9, 209. doi: doi:10.3390/agronomy9040209

Bates, D., Mächler, M., Bolker, B., & Walker, S. (2015). Fitting linear mixed-effects models using lme4. Journal of Statistical Software, 67, 1–48. doi:10.18637/jss.v067.i01

Canci, H., & Toker, C. (2009). Evaluation of yield criteria for drought and heat resistance in chickpea (Cicer arietinum L.). Journal of Agronomy and Crop Science, 195(1), 47–54. doi:10.1111/j.1439-037x.2008.00345.x

Cernay, C., Ben-Ari, T., Pelzer, E., Meynard, J. M., & Makowski, D. (2015). Estimating variability in grain legume yields across Europe and the Americas. Scientific Reports, 5, 11171. doi:10.1038/srep11171

Crépon, K., Marget, P., Peyronnet, C., Carrouée, B., Arese, P. & Duc, G. (2010). Nutritional value of faba bean (Vicia faba L.) seeds for food and feed. Field Crops Research 115:329–339. doi: 10.1016/j.fcr.2009.09.016

de Mendiburu, F. (2010). Agricolae: Statistical Procedures for Agricultural Research. R package version. 1. 1–8.

De Visser, C.L.M., Schreuder, R. & Stoddard, F.L. (2014). The EU’s dependency on soya bean import for the animal feed industry and potential for EU produced alternatives. OCL 21: D407. doi: 10.1051/ocl/2014021

Duc, G., Aleksic, J.M., Marget, P., Mikic, A., Paull, J., Redden, R.J., Sass, O., Stoddard, F.L., Vandenberg, A., Vishniakova, M. & Torres, A.M. (2015). Faba bean. Chapter 5, pp. 141–178 in: Grain Legumes, ed. A.M. de Ron. Springer-Verlag, Berlin, Germany. ISBN 978-1-4939-2796-8 (printed) 978-1-4939-2797-5 (eBook).

FAOSTAT (2019, August 28). United Nations Food and Agriculture Organisation. Retrieved from http://www.fao.org/faostat/en/.

Finlay, K. W., & Wilkinson, G. N. (1963). The analysis of adaptation in a plant-breeding programme. Australian Journal of Agricultural Research, 14, 742–754. doi:10.1071/ar9630742

Frimpong, A., Sinha, A., Tar’an, B., Warkentin, T. D., Gossen, B. D. and Chibbar, R. N. (2009), Genotype and growing environment influence chickpea (*Cicer arietinum* L.) seed composition. J. Sci. Food Agric., 89: 2052–2063. doi:10.1002/jsfa.3690

Jha, A. B., Arganosa, G., Tar’an, B., Diederichsen, A., & Warkentin, T. D. (2012). Characterization of 169 diverse pea germplasm accessions for agronomic performance, Mycosphaerella blight resistance and nutritional profile. Genetic Resources and Crop Evolution, 60, 747–761. doi:10.1007/s10722-012-9871-1

Link, W., Abdelmula, A. A., von Kittlitz, E., Bruns, S., Riemer, H., & Stelling, D. (1999). Genotypic variation for drought tolerance in *Vicia faba*. Plant Breeding, 118, 477–483. doi:10.1046/j.1439-0523.1999.00412.x

Partanen, K., Siljander-Rasi, H., & Alaviuhkola, T. (2006). Feeding weaned piglets and growing-finishing pigs with diets based on mainly home-grown organic feedstuffs. Agricultural and Food Science, 15, 89. doi:10.2137/145960606778644502

Prohens, J., Nuez, F., & Carena, M. J. (2008). Handbook of plant breeding. New York: Springer.

R Core Team (2018). R: A language and environment for statistical computing. R Foundation for Statistical Computing, Vienna, Austria. URL https://www.R-project.org/.

Reckling, M., Döring, T.F., Bergkvist, G., Stoddard, F.L., Watson, C.A., Seddig, S., Chmielewski, F.-M. & Bachinger, J. (2018a). Grain legume yields are as stable as other crops in long-term experiments across northern Europe. Agronomy for Sustainable Development 38: 63. Doi: 10.1007/s13593-018-0541-3

Reckling, M., Döring, T.F., Bergkvist, G., Chmielewski, F.-M., Stoddard, F.L., Watson, C.A., Seddig, S. & Bachinger, J., (2018b). Grain legume yield instability has increased over 60 years in long-term field experiments as measured by a scale-adjusted coefficient of variation. Aspects of Applied Biology 138, Advances in Legume Practice, pp.15–20.

Singmann, H., Bolker, B.,Westfall, J., Aust, F. and Ben-Shachar, M. S. (2019). afex: Analysis of Factorial Experiments. R package version 0.24-1. https://CRAN.R-project.org/package=afex

Stoddard, F. L., Hovinen, S., Kontturi, M., Lindström, K., & Nykänen, A. (2009). Legumes in Finnish agriculture: history, present status and future prospects. Agricultural and Food Science 18:191–205. doi: 10.2137/145960609790059578

Temesgen, T., Keneni, G., Sefera, T., & Jarso, M. (2015). Yield stability and relationships among stability parameters in faba bean (*Vicia faba* L.) genotypes. The Crop Journal, 3, 258–268. doi: 10.1016/j.cj.2015.03.004

Toker, C. (2004). Estimates of broad◻sense heritability for seed yield and yield criteria in faba bean (*Vicia faba* L.). Hereditas, 140, 222–225. doi:10.1111/j.1601-5223.2004.01780.x

